# N-terminal cardiac myosin-binding protein C interactions with myosin and actin filaments using time-resolved FRET

**DOI:** 10.1101/2022.09.07.507024

**Authors:** Fiona L. Wong, Thomas A. Bunch, Victoria C. Lepak, Brett A. Colson

**Affiliations:** Department of Cellular and Molecular Medicine, University of Arizona

**Keywords:** Cardiac muscle, sarcomeric proteins, time-resolved fluorescence resonance energy transfer (TR-FRET), hypertrophic cardiomyopathy (HCM), cMyBP-C

## Abstract

Myosin binding protein-C (cMyBP-C) is a sarcomeric protein responsible for normal contraction and relaxation of the heart. We have used time-resolved fluorescence resonance energy transfer (TR-FRET) to resolve the interactions of cardiac myosin and F-actin with cMyBP-C, focusing on the N-terminal region. The results imply roles of these bound protein complexes in myocardial contraction, with particular relevance to β-adrenergic signaling, heart failure and hypertrophic cardiomyopathy (HCM). N-terminal cMyBP-C domains C0 through C2 (C0-C2) contain binding regions for interactions with both thick (myosin) and thin (actin) filaments. Phosphorylation by protein kinase A (PKA) in the cMyBP-C motif (M-domain) regulates these binding interactions. Our spectroscopic assays detect distances between pairs of site-directed probes on cMyBP-C and either myosin or actin. We engineered intermolecular pairs of labeling sites between donor-labeled myosin regulatory light chain (V105C) or F-actin (C374) and cMyBP-C (S85C in C0, C249 in C1, or P330C in M-domain) to detect interactions. Phosphorylation reduced the interaction of cMyBP-C to both myosin and actin. Further insight was gained from evaluating cMyBP-C HCM mutations T59A, R282W, E334K, and L349R, which revealed increases in myosin-FRET, increases or decreases in actin-FRET, and perturbations of phosphorylation effects. These findings elucidate binding of cMyBP-C to myosin or actin under physiological and pathological conditions, providing new molecular insight into the modulatory role of these protein-protein interactions in cardiac muscle contractility. Further, these findings suggest that the TR-FRET assays are suitable for rapid and accurate determination of quantitative binding for screening physiological conditions and compounds that affect cMyBP-C interactions with myosin or F-actin for therapeutic discovery.

**Significance Statement:** Hypertrophic cardiomyopathy (HCM) is a heritable heart disease involving mutations in genes encoding cardiac muscle proteins. Investigating the underlying molecular mechanisms of HCM mutations provides critical insight into the clinical outcomes and can translate into life-saving therapies. A leading cause of inherited HCM are mutations found in cardiac myosin binding protein-C (cMyBP-C), which binds to both myosin and actin to finely-tune contractility. Efforts in elucidating the details of cMyBP-C interactions with myosin and actin have been limited due to standard techniques that are low-throughput and labor-intensive. We have developed a set of Time-Resolved Fluorescence Resonance Energy Transfer (TR-FRET) assays that report the phosphorylation-sensitive binding of N-terminal cMyBP-C to myosin or actin in a high-throughput plate reader format. We detect altered binding due to phosphorylation and unique changes in HCM mutant cMyBP-C binding to myosin versus actin. Our results are informative for developing precision medicine screening assays and new therapies for HCM.

## Introduction

Hypertrophic cardiomyopathy (HCM) is a heritable cardiac disease that affects one in 200-500 people (1, 2). HCM is characterized by a hypercontractile phenotype and modified protein interactions arising from mutations in sarcomeric proteins, resulting in diastolic dysfunction and cardiac remodeling, which can lead to heart failure (HF). The leading cause of HCM is mutations in the gene encoding cardiac myosin binding protein-C (cMyBP-C), *MYBPC3*, comprising ∼40% of all HCM associated mutations (3). cMyBP-C is a sarcomeric protein responsible for normal contraction and relaxation of the heart. Phosphorylation of cMyBP-C fine-tunes cardiac function by regulating its interactions with myosin and actin (4, 5). Therefore, understanding how normal and HCM mutant cMyBP-C are affected by phosphorylation-sensitive binding to myosin and actin will provide therapeutic insights for health and disease as cMyBP-C has been identified as a target for drug compounds that can restore its proper function to treat HCM and/or HF (6, 7).

cMyBP-C is a ∼140 kDa thick filament-associated protein comprised of 11 immunoglobulin-like and fibronectin-like domains termed C0-C10. The C-terminus (C8-C10) is anchored to the thick filament backbone and the N-terminus (C0-C2) extends to binds to myosin and actin in the unphosphorylated state. The main regulator of cMyBP-C function during force development is phosphorylation of the cMyBP-C motif, known as the M-domain. It connects domains C1-C2 and contains four serine residues that can be phosphorylated: Ser275, Ser284, Ser304, and Ser11 (8, 9). Stimulation of the β-adrenergic fight-or-flight response activates PKA to phosphorylate these phosphoserines in M-domain. In the unphosphorylated state, cMyBP-C acts a brake by sequestering myosin heads into an inhibited, super-relaxed state (SRX) (10), and by decreasing myosin ATPase activity (11), which reduces cross-bridge formation and force generation. At the same time, it is thought that unphosphorylated cMyBP-C activates the thin filament by displacing tropomyosin for myosin-binding without the requirement of Ca^2+^ binding to troponin (12-14). The N-terminal cMyBP-C domains C1-C2 have been shown to interact with the proximal 129 residues of myosin’s subfragment-2 (ΔS2) (15) and C0 was found to interact with myosin regulatory light chain (RLC) (16).

cMyBP-C phosphorylation is commonly dysregulated and reduced in patients with HCM or HF, resulting in dyssynchronous myosin and actin interactions (17). Functional studies suggest that phosphorylated cMyBP-C is cardioprotective whereas the effects of HCM mutations on N-terminal cMyBP-C binding to myosin or actin and upon phosphorylation remain largely unknown. Mice expressing phosphomimetic charge substitutions at protein kinase A (PKA)-mediated phosphorylation sites display preserved cardiac morphology (18), enhanced myocardial relaxation (19), and attenuated age-related cardiac dysfunction (20). It is thought that cMyBP-C undergoes structural changes (21-23) upon PKA-mediated phosphorylation and alters interactions with myosin and actin (24), leading to reduced binding. However, the effect of HCM mutations on phosphorylation-regulated cMyBP-C binding to these regions remains unknown.

We recently developed robust high-throughput fluorescence lifetime-based assays to study the dynamic structure of cMyBP-C (23) and the binding interactions between cMyBP-C and F-actin in normal and diseased states (24). Using one of these assays we have screened 1,280 pharmacologically active compounds and have identified the first three drugs that bind to cMyBP-C and abolish its interactions with F-actin (25). We further characterized these drugs effects on C0-C2-actin binding using a TR-FRET assay (25) that we developed further in this research. In this TR-FRET assay a donor fluorescent probe (Fluorescein-5-maleimide (FMAL)) was attached to actin (on Cys374) and an acceptor probe (tetramethylrhodamine (TMR)) was attached to the N-terminal C0-C2 domain. Binding of C0-C2 to actin brings the acceptor close enough to the donor to result in resonance energy transfer from donor to acceptor probes. This was detected as a change in lifetime (the time it takes for the donor intensity to decay from its maximum following excitation, to ∼37% (1/e)) in the FLTPR (25). Here, we further develop the cMyBP-C actin interaction TR-FRET assays and, for the first time establish an assay to accurately measure interactions between cMyBP-C and myosin. For myosin labeling, the donor, FMAL, was placed on the RLC myosin subunit containing the V105C mutation. RLC has no endogenous cysteine residues and introduction of a cysteine at residue 105 (V105C) provides a convenient site for thiol labeling (26). This was then exchanged onto purified full-length porcine ventricle myosin as described (27). C0-C2 containing a single cysteine in the C0 (S85C), C1 (Cys249), and M (P330C) domains were used for acceptor (TMR) labeling. The FMAL-TMR FRET pair has an R_0_ of 5.5 nm, meaning that probes separated by up to 8.25 nM (∼1.5x R_0_) can be reliably detected. This is an appropriate distance separating the head-neck region of myosin and N-terminal cMyBP-C domains.

Using these FRET assays, we studied unphosphorylated and phosphorylated C0-C2 binding to myosin and actin. We then examined the effects of 5 potential C0-C2 HCM mutations on this binding. Three of the mutants (E334K, L349R, and L352P) were chosen due their localization in the tri-helix bundle within the motif (24). One of the mutations (R282W) disrupts one of the PKA phosphorylation target sites in the motif, resulting in decreased effects of PKA on actin binding (24). Finally, one of the mutations (T59A in C0) has been reported to influence binding to RLC (16), however alanine is found in this position of C0 in several other species and may simply be a non-pathogenic variant in humans. We previously studied the actin binding properties of three of these mutations (R282W, E334K, and L352P) using an environmentally-sensitive single-probe fluorescent lifetime assay (i.e., time-resolved fluorescence, TR-F) (24) and others have investigated mutation effects on myosin/actin binding or phosphorylation (21-23).

Finally, we have tested our new assays for their usefulness in future high throughput screens for therapeutic compounds that modulate C0-C2 actin and myosin binding.

## Results

### TR-FRET-based techniques to detect the interaction of N-terminal C0-C2 and myosin

Site-directed mutagenesis introduced a cysteine in the RLC subunit at residue 105, V105C (Fig. 1B). To ensure that the FMAL on one RLC did not interact with FMAL on a neighboring RLC we exchanged 10% labeled RLC onto myosin. This was achieved by using a mixture of FMAL-labeled and unlabeled RLC for exchange (27). RLC exchange was monitored by analyzing SDS-PAGE gels for fluorescence and protein staining (Fig. 2A). F-actin was labeled at site Cys-374 (Fig. 1C). Labeled and unlabeled G-actin were also mixed to achieve 10% labeled monomers in the final F-actin filaments. To monitor the myosin-binding and actin-binding, we inserted acceptor probes at different positions (C0; Cys85, C1; Cys249; the tri-helix bundle in the motif; Cys330) in the cMyBP-C N-terminal C0-C2 (Fig. 1E-1G). C0-C2 contains 5 endogenous cysteines, which were strategically removed to allow for introduction of single cysteines (Fig. 1F, 1H) (23). Since Cys249 of the C1 domain is an endogenous, surface-accessible thiol, the remaining 4 Cys were removed to generate this protein (Fig. 1G).

**Fig. 1.**
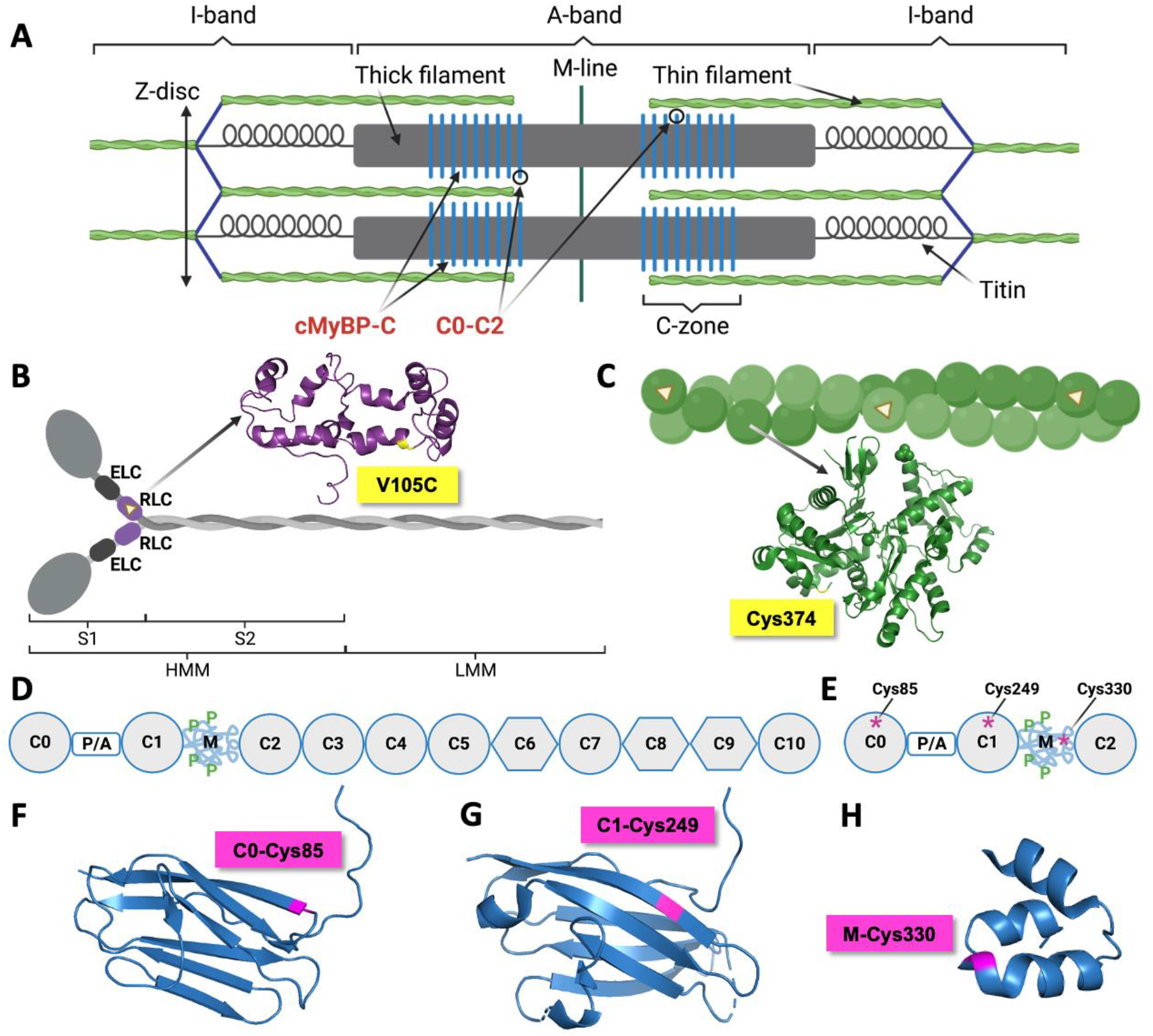
Organization of the sarcomere and the positioning of the TR-FRET probes on myosin, actin, and cMyBP-C. (A) The sarcomere is the smallest functional unit of the muscle, which spans from Z-disc to Z-disc. It is comprised of the actin thin filament (I-band) and myosin thick filament (A-band), which slides across one another to generate proper contraction and relaxation of the muscle. The C-zone is the thick filament region where cMyBP-C molecules are located (blue vertical stripes). The sarcomere is stabilized by the titin protein, which anchor in the Z-disc and extend to the M-line. (B) β-cardiac myosin is a hexamer comprised of two heavy chains, two essential light chains (ELC) and two regulatory light chains (RLC). The heavy chains can be proteolyzed into heavy meromyosin (HMM) and light meromyosin (LMM). LMM is the distal portion of the heavy chain, which mediates myosin filament assembly. HMM contains Subfragment-1 (S1), the motor and catalytic domain of myosin, and Subfragment-2 (S2), the first 126 amino acid residues of the **α**-helical coiled-coil tail. ∼10% of myosin is labeled with FMAL, the donor probe, on the RLC subunits (purple ovals) at site V105C (PDB: 5TBY). (C) F-actin (green circles) is a filamentous polymer consisting of G-actin monomers and is similarly labeled with ∼10% FMAL (yellow triangles) on site Cys374 (PDB: 3HBT). (D) Full-length cMyBP-C contain domains C0-C10. Immunoglobulin-like domains are shown as circles and fibronectin type-III domains are shown as hexagons. The C-terminus (C8-C10) is anchored to the thick filament backbone and the N-terminus (C0-C2) binds to myosin and actin. (E) N-terminal domains C0 through C2 (C0-C2) contains the proline/alanine-rich linker (P/A) and the M-domain (M) that contains phosphorylation sites (P). Acceptor probe TMR is inserted in the C0 (Cys85), C1 (Cys249), and M (Cys330)-domain. (F) C0-domain structure (cyan, PDB: 2K1M) with TMR (pink) on site Cys85. (G) C1-domain structure (PDB: 2V6H) with TMR on site Cys249. (H) C0-domain structure (PDB: 2LHU) with TMR on site Cys330.

**Fig. 2.**
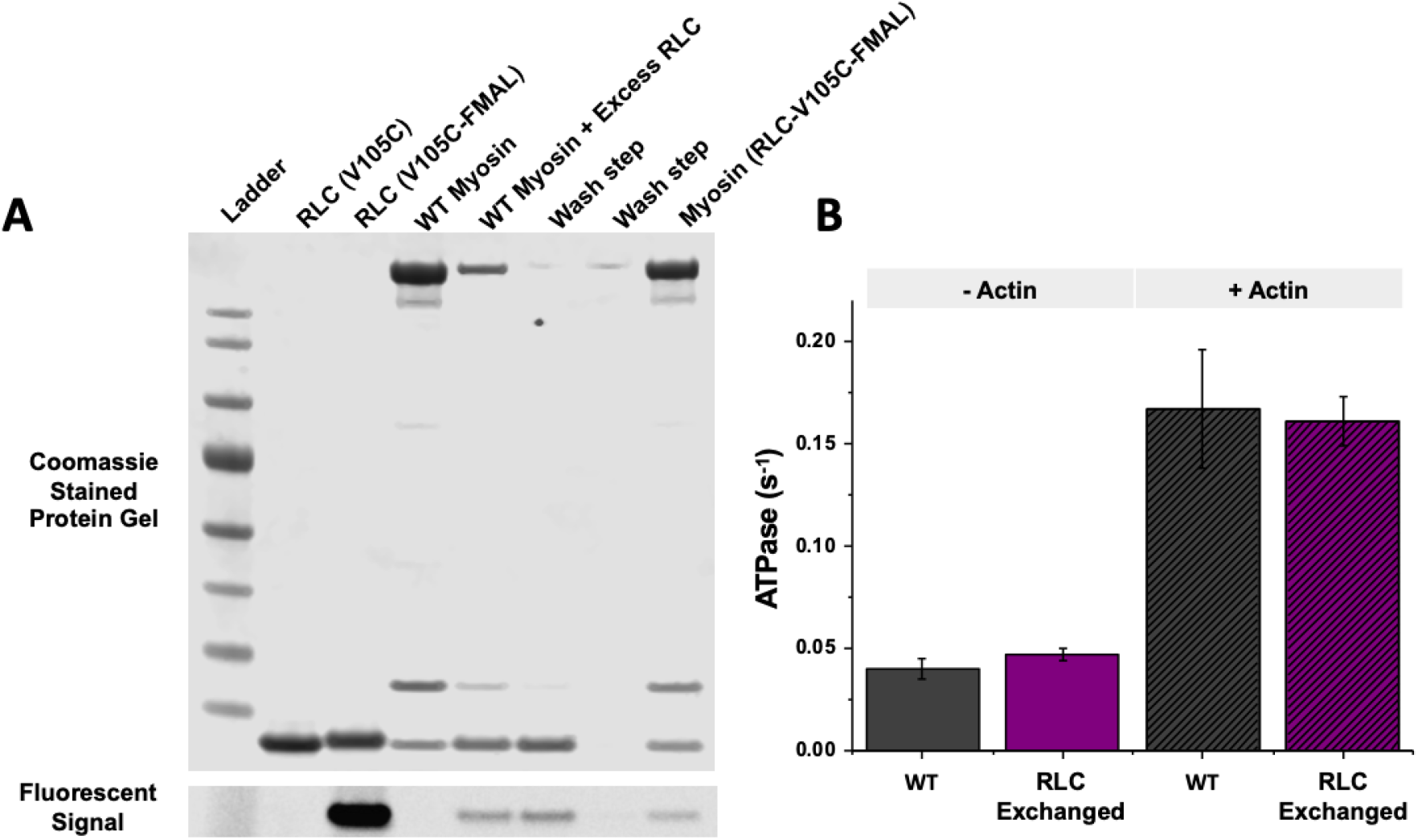
Donor-labeled RLC exchange onto WT full-length purified myosin method and ATPase activity. (A) Representative Coomassie stained SDS-PAGE gel demonstrates each step of the Myosin-RLC exchange. The exchange mixture contains 25 μM RLC with 5 μM of myosin. The 25 μM RLC contains 10% FMAL-labeled RLC-V105C. A series of wash and high-speed centrifugation steps are performed to remove unincorporated RLC. The final myosin product contains ∼10% FMAL on site RLC-V105C, which prevents intramolecular FRET between the neighboring RLC molecules on myosin. Further details can be found in the methods. 1.5 μg of RLC or ∼5 μg of myosin was loaded per lane. FMAL-labeled RLC and its incorporation into myosin can be traced by the band present in the fluorescent signal. (B) The steady-state ATPase assay was used to verify whether the donor-labeled RLC-exchanged myosin has retained normal function. The ATPase activity of 1 μM of WT myosin (0.040 ± 0.005) and RLC-exchanged myosin (0.047 ± 0.003) were similar. Actin-activated ATPase activity (+4 μM of actin) of the WT myosin (0.167 ± 0.029) and RLC-exchanged myosin (0.161 ± 0.012) were also comparable. In conclusion, the function of donor-labeled myosin was not altered following the RLC exchange. Data shown as mean ± S.E., n = 4-6 (3-4 preparations).

To ensure that the FMAL-RLC exchanged myosin maintained the same functionality as non-exchanged myosin with native RLC, we utilized the steady-state ATPase assay to test its function compared to this wild-type control (WT) (Fig. 2B). Using a regenerative system with enzymes (pyruvate kinase and lactate dehydrogenase) that depletes NADH as ATP is hydrolyzed by myosin, we measured the colorimetric decrease in NADH and calculated the rate of hydrolysis of one ATP per catalytic cycle of myosin. The basal and actin-activated myosin ATPase activity of 0.040 ± 0.005 s^-1^ and 0.167 ± 0.029 s^-1^ for the RLC-exchanged myosin was not different from control myosin with ATPase activity of 0.047 ± 0.003 s^-1^ and 0.161 ± 0.012 s^-1^, respectively. These results demonstrated that the RLC exchanged myosin was unaffected by the exchange procedure with the V105C substitution and retained normal ATPase function.

### C0-C2 phosphorylation-dependent changes in FRET to myosin and actin

TR-FRET waveform decays of FMAL-labeled myosin and actin in the presence of TMR-labeled C0-C2 were analyzed by one-exponential fitting (see Methods) to determine fluorescence lifetime (Fig. 3). FMAL lifetime decreased for both myosin and actin in the presence of TMR-C0-C2, indicative of binding between C0-C2–myosin (Fig. 2A) and C0-C2–actin (Fig. 2D). This effect was reduced with phosphorylated C0-C2, consistent with reduced binding (Fig. 2B, 2E). Myosin and Actin TR-FRET assays were fit to binding curves for each TMR-C0-C2 (with the acceptor probes on Cys85, Cys249, and Cys330). Analysis of results with 1 μM FMAL-myosin or 1 uM. FMAL-actin revealed that PKA phosphorylation of TMR-C0-C2 significantly reduced FRET Efficiency in both assays (Fig. 4). The K_D_, maximum FRET (FRET_max_), and adjusted R^2^ (adj. R^2^) was determined by fitting the data to a quadratic model (Table S1). Reduced FRET due to phosphorylated C0-C2 in the presence of actin maintains the shape of a binding curve. The C0-C2–actin binding curves presented (Fig. 4D-F) had an adjusted R^2^ of 0.99-1.00, which shows how well the data points fit to the model. However, the adjusted R^2^ of the phosphorylated C0-C2-myosin were significantly lower at 0.60-0.88 (Table S1). Due to the poor fit, myosin TR-FRET with phosphorylated C0-C2 may either be reflecting decreased or completely abolished binding. Importantly, FRET decreased in the presence of PKA phosphorylation of C0-C2 with myosin and actin.

**Fig. 3.**
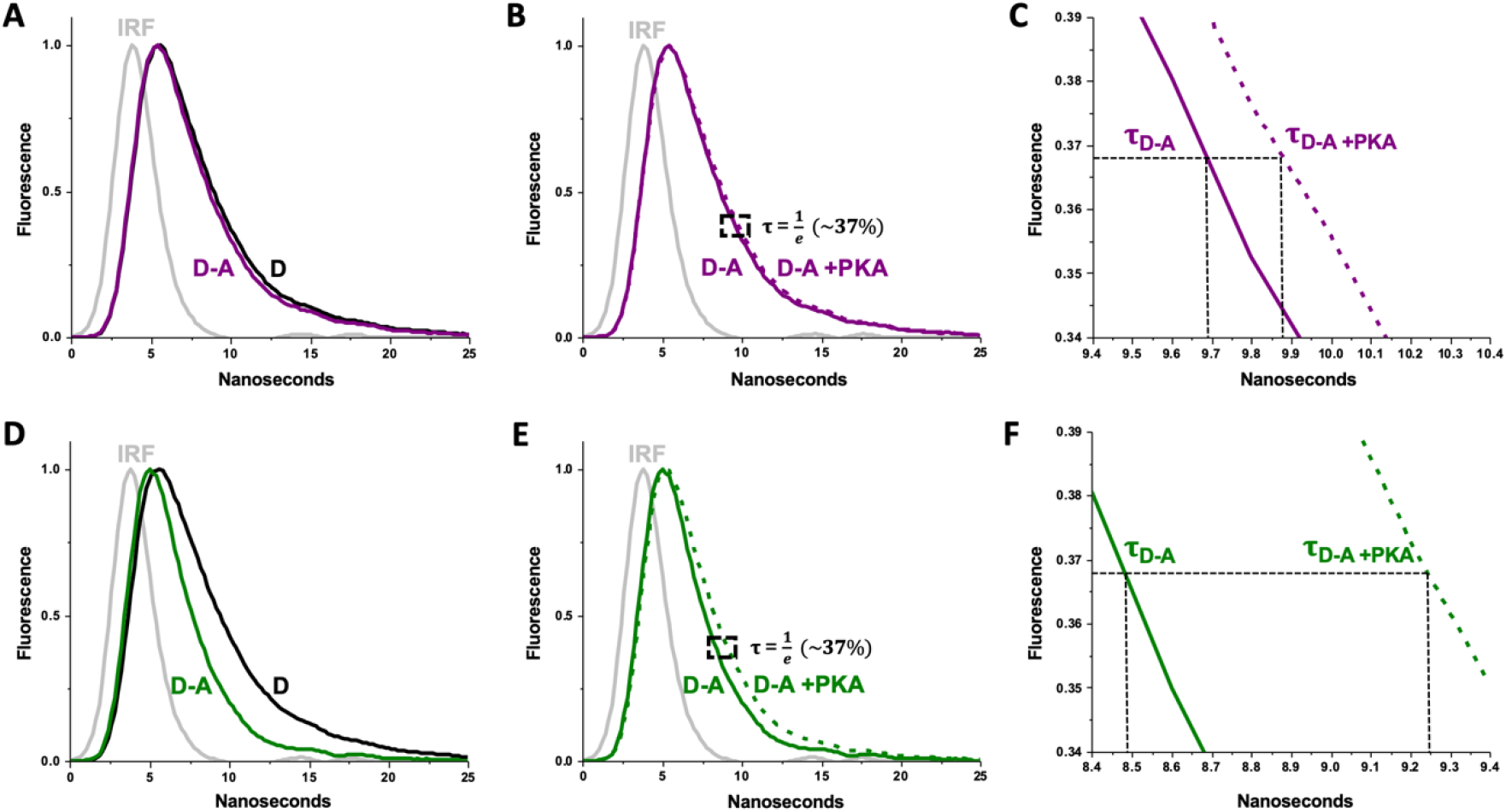
Fluorescence lifetime decays for TR-FRET of FMAL-labeled myosin and actin with TMR-labeled C0-C2. (A) Normalized TR-FRET waveform decays of 1 μM of myosin and 20 μM C0-C2. Instrument response function (IRF) of water scattering is shown in grey. Donor-only (D, black) and donor-acceptor (D-A, purple) samples exhibit waveform decays after initial rising phase due to FMAL excitation. (B) Waveform decays of D-A labeled unphosphorylated C0-C2 (purple) and PKA-phosphorylated C0-C2 (dotted purple). Region of fluorescence lifetime, where **τ** decays to 1/e (∼37%) of peak intensity, is highlighted by dotted box. (C) Magnified dotted box region in panel B. (D) Normalized TR-FRET waveform decays of 2 μM of actin and 10 μM C0-C2. Donor-only (D, black) and donor-acceptor (D-A, green) samples. (E) D-A labeled unphosphorylated C0-C2 (green) and PKA-phosphorylated C0-C2 (dotted green). The effect of PKA phosphorylation of C0-C2 is more pronounced on actin than on myosin. (F) Magnified dotted box region in panel E.

**Fig. 4.**
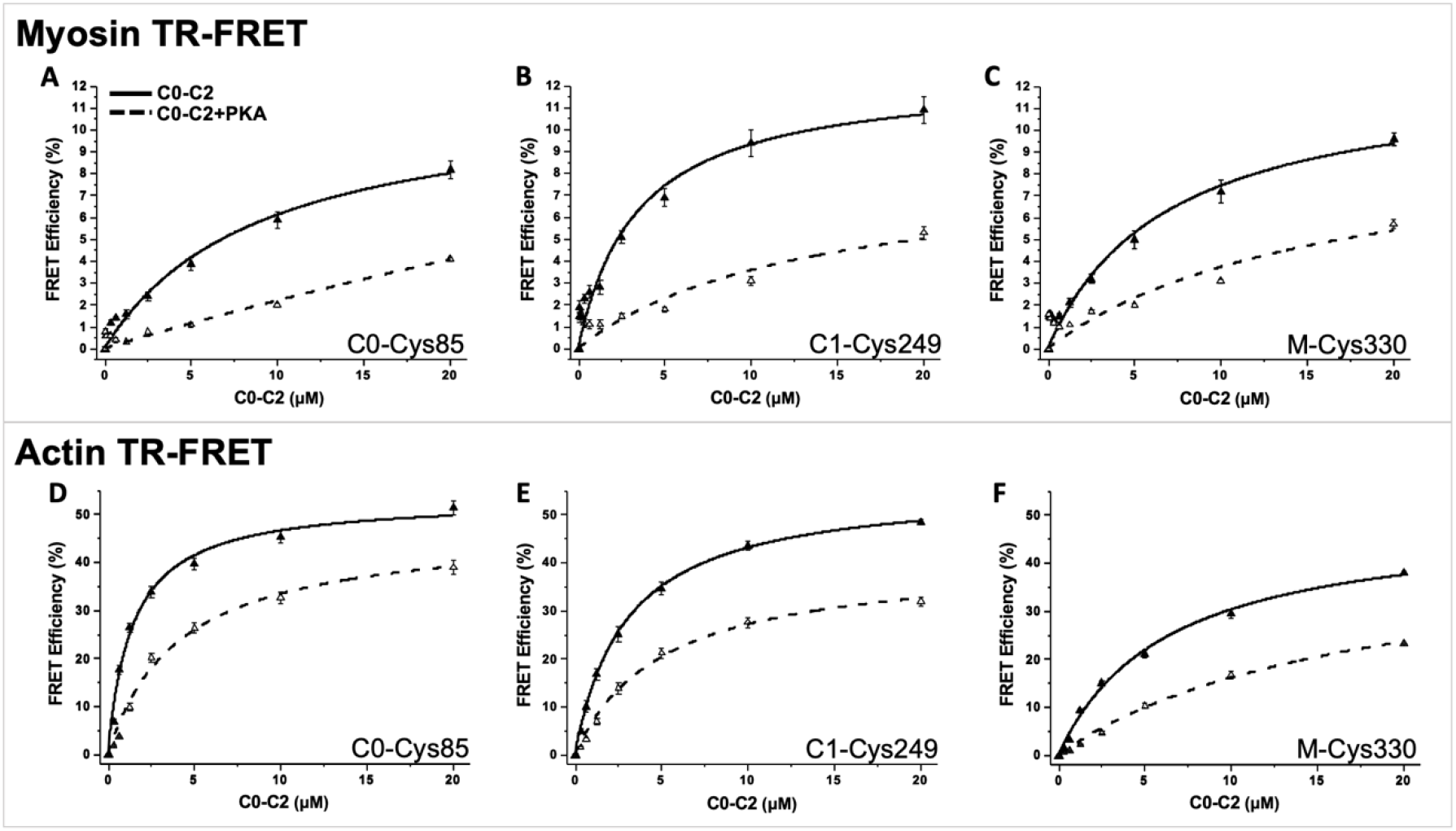
TR-FRET binding assays for myosin and actin using three different acceptor-probe sites on C0-C2. Unphosphorylated C0-C2 (solid lines) and phosphorylated C0-C2 (dotted lines) were assayed. (A-C) TR-FRET using FMAL-myosin. (D-F) TR-FRET using FMAL-actin. (A and D) Acceptor is located on C0-domain at site Cys85. (B and E) Acceptor is located on C1-domain at site Cys249. (C and F) Acceptor is located on M-domain at site Cys330. Refer to supplemental Table 1 for binding affinities (K_D_), maximal FRET Efficiencies (FRET_MAX_), and adjusted R^2^. Data are provided as mean ± SE (N=3, n=10-15).

### C0-C2 HCM mutation effects on myosin and actin TR-FRET

We used these new assays to test the effects of 5 potential HCM mutations, one in C0, one in a PKA phosphorylation target sequence and 3 in the tri-helix bundle located in the M-domain (Fig. 5). All mutations were tested with myosin and actin TR-FRET and all of the mutants were tested in C0-C2 separately labeled at each of the 3 acceptor probe sites. In Table 1, we looked at the relative FRET of 2.5 μM C0-C2 bearing each mutation compared to the same concentration of wild type protein interacting with myosin or actin. From the binding curves (Fig. 4), 2.5 μM C0-C2 shows clear binding effects that are submaximal. At this concentration we expect to be able to detect both increases and decreases in interactions using these FRET assays. This is further verified for all mutants, both without and with phosphorylation, by FRET observed with labeled myosin at 5 and 10 μM C0-C2 (Fig. S2). For actin, binding curves for all mutants (Fig. S3) illustrate the usefulness of 2.5 μM C0-C2 for comparisons given in Table 1.

**Fig. 5.**
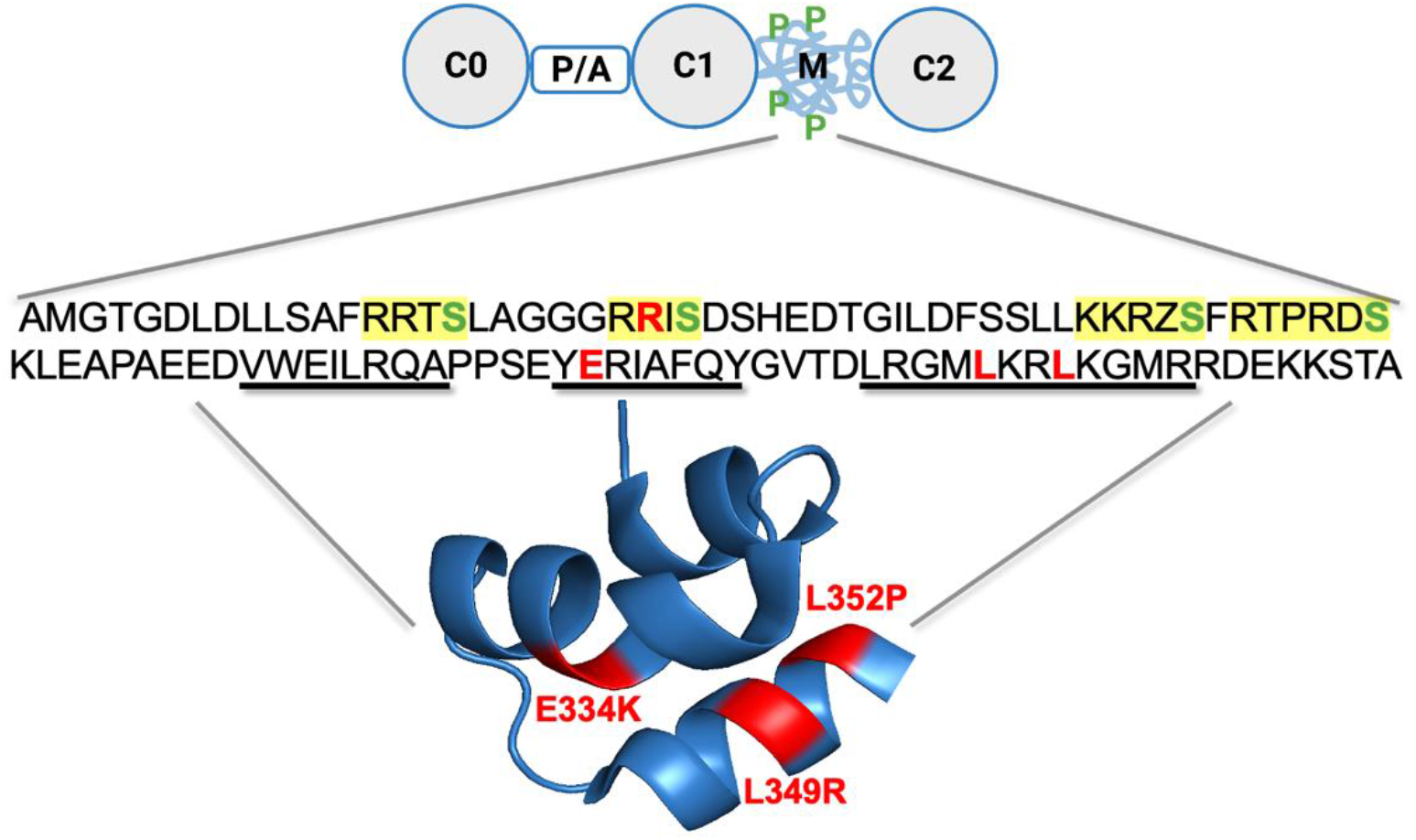
cMyBP-C organization and HCM mutants in C0-C2 tested. C0-C2 domains containing P/A linker and M-domain containing phosphorylation sites are shown. Sequence of M-domain and locations of HCM mutations tested for binding. PKA phosphorylatable Ser residues are in green, PKA recognition sequences are highlighted in yellow, and HCM mutations are in red. Helix residues in the tri-helix bundle are indicated with thick underlines. Tri-helix bundle (PDB: 5K6P) contains E334K, L349R, and L352P mutations. R282W mutation is location near a phosphorylation recognition site. Mutation T59A is not seen here as it is in the C0-domain.

**Table 1.** Binding effects of HCM mutants in C0-C2 to myosin and actin relative to WT FRET at 2.5 μM.

| **C0-C2** | **PKA** | **Myosin** |  |  | **Actin** |  |  | **Relative to WT FRET at 2.5 μM** |
| --- | --- | --- | --- | --- | --- | --- | --- | --- |
| **Acceptor Site** |  | **C0** | **C1** | **M** | **C0** | **C1** | **M** | **<0.5** |
| **T59A** | - | 0.91 | 1.15 | 0.96 | 0.97 | 0.92 | 1.21 | 0.8-1.2 |
|  | + | 0.80 | 1.04 | 1.24 | 0.89 | 0.96 | 0.98 | 1.2-1.5 |
| **R282W** | - | 1.01 | 0.98 | 1.02 | 1.03 | 0.97 | 1.10 | 1.5-2.0 |
|  | + | 1.51 | 1.24 | 1.64 | 1.37 | 1.37 | 2.05 | >2.0 |
| **E334K** | - | 1.71 | 1.71 | 1.96 | 1.18 | 1.08 | 1.97 |  |
|  | + | 1.67 | 1.70 | 2.00 | 0.45 | 1.05 | 2.03 |  |
| **L349R** | - | 1.45 | 1.72 | 1.60 | 1.22 | 1.10 | 1.89 |  |
|  | + | 1.45 | 1.43 | 1.71 | 1.04 | 1.31 | 2.66 |  |
| **L352P** | - | 1.46 | 1.25 | 1.54 | 1.36 | 1.21 | 1.31 |  |
|  | + | 1.36 | 0.99 | 2.01 | 2.27 | 1.74 | 2.60 |  |

Mutations T59A, R282W, E334K, L349R, and L352P in C0-C2 with an acceptor site in C0 (S85C), C1 (Cys249), or M-domain (P330C) were tested with 1 μM of myosin or actin. The relative FRET value of each HCM mutation at 2.5 μM to WT C0-C2 at 2.5 μM was calculated to evaluate the respective changes for each probe site. Red indicates <50% of a decrease relative to WT, grey indicates an 80% to 120% change (increase or decrease), light green indicates a 120% to 150% increase, green indicates a 150% to 200% increase, and dark green indicates >200% of an increase.

**T59A**, in C0, at 2.5 μM C0-C2, displayed FRET values that were within 20% of that found for wild type C0-C2 in 5 of the 6 conditions tested for both actin and myosin (3 probe acceptor sites, minus and plus phosphorylation). On myosin the probe in the tri-helix bundle (Cys330) did show a 24% increase in FRET with myosin when T59A was phosphorylated and a 21% increase in FRET with actin in the unphosphorylated state (Table 1). **R282W**, in a PKA target sequence, altered phosphorylation and actin interactions (23, 24). Our new FRET assays found no effects on the binding of unphosphorylated R282W to either myosin or actin (FRET was within 10% of wild type for all 3 FRET acceptor probes). The phosphorylated forms of R282W all displayed increased FRET with both myosin (24%-64% increases) and actin (37%-105% increases). This is consistent with a reduction in phosphorylation and our previous results (24). The largest effects on FRET for phosphorylated R282W with both myosin (64%) and actin (105%) were observed when the acceptor probe was located in the tri-helix bundle at Cys330. **E334K**, in the tri-helix bundle, increased myosin FRET in all 6 conditions binding by 66%-100%. With actin, probes on C0 and C1 showed little change (within 18% of wild type) or a significant decrease to 45% of wild type FRET in the case of the C0 probe when phosphorylated. When the probe is in the tri-helix bundle E334K, unphosphorylated and phosphorylated, promoted a doubling of the FRET (Table 1). **L349R** consistently increases myosin FRET by 43%-72% in all conditions tested. Actin FRET with L349R gave mixed results with probes on C0 and C1. Minimal change in phosphorylated C0 and unphosphorylated C1 (4% and 10% respectively). Moderated changes were observed in unphosphorylated C0 and phosphorylated C1 (22% and 31% respectively). The probe on the tri-helix bundle responded strongly to L349R with increase of 89% (unphosphorylated) and 166% (phosphorylated). **L352P** increases both myosin and actin FRET. With both myosin and actin, the effects were largest for the probe in the tri-helix bundle, followed by the probe in C0 and then the one in C1. On actin, large effects (74%-160%) were observed for all probe sites when L352P was phosphorylated. For myosin, the phosphorylated form showed a large effect (101% increase) on the probe in the tri-helix bundle, a moderate effect (36% increase) on the probe in C0 and no effect on the probe in C1 (Table 1).

### Assessing high-throughput screening (HTS) assay suitability of myosin and actin in the presence and absence of C0-C2 binding

We tested the suitability of the myosin and actin TR-FRET assays for suitability in HTS. FRET measured in the presence and absence of C0-C2 binding represents a mock screen for identification of small-molecule drugs that inhibit C0-C2 binding to myosin or actin. These FRET values were used to determining a *Z′*-factor (see Materials and Methods for details). Lifetime averages and standard deviations (S.D.) of the donor probe (FMAL) in acceptor (TMR) labeled C0-C2 (Cys249) were determined in the presence and absence of C0-C2. These values were then used to calculate *Z′*-factor (Eq. 3). *Z′*-factors less than 0 indicate that screens done under the conditions tested would be useless in HTS. *Z′*-factors between 0.0 and 0.5 indicate a usable screen and *Z′*-factors between 0.5 and 1.0 are indicative of an excellent HTS assay (28). Using a 384-well plate, FMAL-myosin and FMAL-actin were tested for TMR-C0-C2 (Cys249) binding-induced changes in lifetime. Table 2 shows 4 independent preparations with 3-5 wells with and without C0-C2. The *Z′*-factor ranges from 0.76-0.90 for myosin and from 0.94-0.95 for actin, which classifies our assays as “excellent” for use in HTS for compounds modulating cMyBP-C interactions with myosin or actin.

**Table 2.** *Z′*-factor test comparing the absence and presence of C0-C2 lifetimes.

| **Myosin** | **Prep** | **C0-C2** | **Lifetimes (ns)** |  |  |  |  | **AVG** | **STD** | **Z'** |
| --- | --- | --- | --- | --- | --- | --- | --- | --- | --- | --- |
|  | 1 | - | 3.66 | 3.66 | 3.67 | 3.67 | 3.67 | **3.67** | **0.005** | **.76** |
|  |  | + | 3.51 | 3.51 | 3.51 | 3.50 | 3.52 | **3.51** | **0.007** |  |
|  | 2 | - | 3.50 | 3.51 | 3.51 | 3.50 | 3.52 | **3.51** | **0.008** | **.84** |
|  |  | + | 3.25 | 3.25 | 3.25 | 3.26 | 3.25 | **3.25** | **0.006** |  |
|  | 3 | - | 3.43 | 3.42 | 3.44 | 3.43 | 3.43 | **3.43** | **0.005** | **.85** |
|  |  | + | 3.24 | 3.25 | 3.24 | 3.24 | 3.24 | **3.24** | **0.004** |  |
|  | 4 | - | 3.79 | 3.78 | 3.80 | 3.80 | 3.80 | **3.79** | **0.005** | **.90** |
|  |  | + | 3.52 | 3.53 | 3.52 | 3.52 | 3.52 | **3.52** | **0.004** |  |
|  | **Prep** | **C0-C2** | **Lifetimes (ns)** |  |  |  |  | **AVG** | **STD** | **Z'** |
| **Actin** | 1 | - | 4.08 | 4.07 | 4.09 | -- | -- | **4.08** | **0.007** | **.94** |
|  |  | + | 2.85 | 2.82 | 2.84 | -- | -- | **2.84** | **0.018** |  |
|  | 2 | - | 4.20 | 4.21 | 4.18 | 4.21 | 4.21 | **4.20** | **0.013** | **.95** |
|  |  | + | 2.64 | 2.65 | 2.63 | 2.62 | 2.64 | **2.64** | **0.011** |  |
|  | 3 | - | 3.81 | 3.80 | 3.80 | 3.80 | 3.79 | **3.80** | **0.007** | **.95** |
|  |  | + | 2.44 | 2.40 | 2.42 | 2.43 | -- | **2.42** | **0.017** |  |
|  | 4 | - | 3.81 | 3.80 | 3.80 | 3.80 | 3.79 | **3.80** | **0.007** | **.95** |
|  |  | + | 2.44 | 2.40 | 2.42 | 2.43 | -- | **2.42** | **0.021** |  |

Lifetimes of 5 $\mu\text{M}$ D-A labeled C0-C2 (Cys249) are shown here with 1 $\mu\text{M}$ of myosin or 2 $\mu\text{M}$ of actin. 4 preps are shown with 3-5 wells ± C0-C2. Z'-factor is defined as the difference between 3x S.D divided by the difference in the mean signal (see Materials and Methods).

## Discussion

Both actin and myosin interactions have been proposed to be important for the mechanism by which cMyBP-C modulates contractility in healthy and diseased myocardium, including PKA-mediated phosphorylation-dependent binding (Fig. 1) (for review, (29-31)). Therefore, we developed a set of assays useful to monitor these interactions that are sensitive variables including binding level, phosphorylation level, and mutations in cMyBP-C that cause HCM disease (Fig. 2-4, Table 1).

Here, we show three versions of cMyBP-C N-terminal fragment C0-C2 containing a FRET acceptor site in either C0, C1, or the tri-helix bundle in the M-domain (Fig. 1 E-H). Each of these probe sites, for both actin and myosin interactions, are responsive to PKA treatment and HCM mutations in C0-C2 (Fig. S2 and S3, Table 1 and S1) (primarily located in M-domain (Fig. 5)). However, each version of C0-C2 shows a different binding curve, suggesting that the different probe sites are monitoring different modes and/or interfaces of actin or myosin binding.

Large changes in intermolecular TR-FRET were detected due to the key variables tested and with low variability in the fluorescence lifetime measurements of technical repeats in wells in the plate reader format (Table 2). Therefore, these assays will be useful for screens and monitoring effects of cMyBP-C mutations and small-molecule drugs.

In the present study, we developed a cMyBP-C–myosin binding assays and extended our cMyBP-C–actin binding assays (24, 25), relevant for physiologic, pathologic, and therapeutic purposes. By utilizing this technology alongside our previously established cMyBP-C–actin binding assay, we monitored interactions of normal and HCM mutant cMyBP-C N-terminal domains C0-C2, in unphosphorylated and phosphorylated forms, to myosin or actin. The validation of these assays makes screening of small-molecule drugs that alter the binding between human C0-C2 and myosin or actin feasible using the TR-FRET technique performed in a FLTPR instrument.

## Materials and Methods

### Myosin filament preparations

Cardiac myosin from porcine ventricles (*Pel Freeze)* was prepared as described by Rohde *et al*. (32) and further purified with ion-exchange chromatography (HiPrep DEAE FF 16/10 column) to remove contamination of thin filament proteins. The column was equilibrated with 40mM NaPyrophosphate, pH 7.5 and the starting myosin was eluted in a gradient of 0-500mM KCl, 20 mM NaPyrophosphate, over 100 mL at 2 mL/min, collecting 5 mL fractions (AKTA Prime Plus, GE). SDS-page was used to pool the fractions with <2% of actin contamination. The final myosin was dialyzed into 600 mM KCl, 25 mM KP_i_, 2 mM DTT, pH 7.0 buffer. For storage at -80°C, 150 mM sucrose was added to the purified myosin and was flash frozen in a dropwise manner in liquid nitrogen.

### RLC preparation and labeling

Human RLC with a single cysteine at position 105 engineered using a Q5 Site-Directed Mutagenesis Kit (New England Bio Labs) was expressed in pET45b vectors encoding E. coli optimized codons (GenScript) and was purified by inclusion body isolation (33). RLC was purified from degraded products by ion exchange chromatography (Hitrap_Q_XL_5mL column). The column was equilibrated with 4 M Urea, 30 mM Tris-HCl, 50 mM NaCl, 1 mM DTT NaCl, pH 7.5. The starting RLC was eluted in a gradient of 0-250mM NaCl over 50 mL at 1 mL/min, collecting 1 mL fractions (AKTA Prime Plus, GE). The RLC myosin was dialyzed into 50 mM Tris, 50 mM NaCl, 40% sucrose, pH 7.5 and was flash frozen in liquid nitrogen and stored at −80°C until use.

Fluorescein-5-maleimide (FMAL) was used to label RLC as the donor probe. RLC (50 μM) was treated with the reducing agent TCEP (200 μM) for 30 min at 23°C while rocking. FMAL was added (stored as 20 mM stock in DMF at -80°C) to a final concentration of 275 μM. Labeling was done for 2 h at 23°C and terminated by the addition DTT (to 5x molar excess of dye concentration to terminate the reaction of unbound dye). Unincorporated dye was removed by extensive dialysis into exchange buffer (50 mM NaCl, 5 mM EDTA, 2 mM EGTA, 10 mM NaPO_4_, pH 7.5). Following the dialysis, labeled RLC was centrifuged for 30 min at 100,000 RPM (350,000 *xg*) in a Beckman TLA-120.2 rotor to remove any precipitated dye and insoluble protein. The degree of labeling was determined by the FMAL’s extinction coefficient.

### RLC exchange onto myosin

The method followed for the RLC exchange onto myosin was described in Kast 2010 *et al*. (27) and was adjusted for the use of full-length myosin. Labeled RLC and purified myosin were dialyzed into the exchange buffer (50 mM NaCl, 5 mM EDTA, 2 mM EGTA, 1 mM DTT, 10 mM NaPO_4_). RLCs were exchanged by adding 25 μM of RLC (2.5 μM FMAL labeled RLC, 22.5 μM unlabeled RLC), 5 μM of myosin, 5 mM EDTA, 2 mM ATP, and 1 mM DTT. After a 50 min incubation at 42°C, 15 mM MgCl_2_ was slowly added to stop the exchange. To remove excess RLC, the mixture was centrifuged at 200,000 *xg* in a Beckman TLA-120.2 rotor, pelleting the myosin filaments. The supernatant containing excess RLC was removed, and the pellet was washed with exchange buffer containing 15 mM MgCl2. The pellet was centrifuged at 200,000 *xg* for an additional 5 min in the wash solution. The supernatant was removed, and the final pellet was resuspended in 600 mM NaCl, 0.2 mM EDTA, 25 mM Tris, pH 7.0. After homogeneous resuspension, the sample was clarified by centrifugation at 4°C, 15,000 rpm (21,000 *xg*) for 10 min in an Eppendorf 5424R benchtop microfuge and was dialyzed into 75-20-3 buffer (75 mM KCl, 20 mM Tris, 3 mM MgCl_2_, pH 7.0) for TR-FRET experiments.

### Actin filament preparations and labeling

Actin was prepared from rabbit skeletal muscle by extracting acetone powder in cold water as described in Bunch *et al*. (24, 34). Actin was labeled as F-actin. First, 50 μM G-actin (in 10 mM Tris, 0.2 mM CaCl_2_, 0.2 mM ATP, pH 7.5) was brought to 20 mM Tris (pH 7.5) and then polymerized by the addition of MgCl_2_ to a final concentration of 2 mM and KCl to a final concentration of 100 mM. Polymerization was allowed to proceed for 1 hour at 23°C. FMAL was added to a final concentration of 0.5 mM (from a 20 mM stock in DMF). Labeling was for 1 h at 23°C and stopped by the addition of a 5-fold molar excess of DTT. Labeled F-actin was collected by centrifugation (30 min, 100,000 rpm (350,000 ×*g*) in a Beckman TLA-120.2 rotor at 4°C). The F-actin pellet was rinsed 3x with actin labeling G-buffer (5 mM Tris, 0.2 mM CaCl_2_, 0.5 mM ATP, pH 7.5) containing 3 mM MgCl_2_ then 1x with actin labeling G-buffer and then resuspended in actin labeling G-buffer. Labeled G-actin was clarified to remove any remaining F-actin or precipitated FMAL by centrifugation (10 min, 90,000 rpm in a Beckman TLA-120.2 rotor at 4°C). The extent of labeling (typically around 70%) was determined by by UV-vis spectrophotometry. Labeled G-actin was mixed with unlabeled G-actin to achieve a 10% labeled mixture. This was polymerized by the addition of MgCl_2_ to a final concentration of 2 mM and KCl to a final concentration of 100 mM and dialyzed against MOPS actin-binding buffer (M-ABB; 100 mM KCl, 10 mM MOPS, pH 6.8, 2 mM MgCl_2_, 0.2 mM CaCl_2_, 0.2 mM ATP, 1 mM dithiothreitol [DTT], and 1 mM sodium azide). Any bundled actin was removed by centrifugation at 4°C, 15,000 rpm (21,000 *xg*) for 10 min in an Eppendorf 5424R benchtop microfuge. Actin concentration and % FMAL labeling was again determined by UV-vis spectrophotometry. The labeled F-actin was stabilized with the addition of phalloidin (to the same concentration as actin).

### Steady-state basal and actin-activated ATPase activity of myosin

NADH-coupled assay was used to measure the steady-state ATPase activity of full-length purified myosin (± RLC exchange) and was performed at 23°C in buffer containing 10 mM KCl, 4 mM MgCl_2_, 20 mM Tris-HCl, and at pH 7.5. Each reaction contained 1 μM of myosin ± 4 μM of actin and 2 mM phosphoenolpyruvate, 0.3 mM NADH, 38 U/ml pyruvate kinase, 50 U/mL lactate dehydrogenase, and 2 mM of MgATP. Changes in NADH absorption due to oxidation at 340 nm was captured by using a Beckman DU730 UV-Vis spectrophotometer for 15 min. The negative slope of the absorbance is proportional to the amount of ATPase activity. The extinction coefficient of NADH (ε_340_ = 6220 M^-1^ cm^-1^) was used to convert the absorbance at 340 nm to [ADP]. A plot of the time course was generated and fit to a linear function to determine the slope of the steady-state ATPase rate in units of [ADP] per unit time. This observed rate was divided by the concentration of myosin in the reaction. The units of the rates are ADP myosin^-1^ s^-1^ but is commonly expressed as s^-1^, such that one ATP is hydrolyzed per catalytic cycle of myosin (35).

### Recombinant human cMyBP-C and labeling

pET45b vectors encoding E. coli optimized codons for the C0-C2 portion of human cMyBP-C with N-terminal 6x His tag and TEV protease cleavage site were obtained from GenScript. C0-C2 mutants were engineered using a Q5 Site-Directed Mutagenesis Kit (New England Bio Labs). Substitution mutations were performed to generate C0-C2 constructs containing a single cysteine located in the C0-domain at position 85, C1-domain at position 249, or M-domain at position 330 (termed C0-C2^S85C^, C0-C2^Cys249^, and C0-C2^P330C^). Five endogenous cysteines were removed to generate C0-C2^Cys85^ and C0-C2^Cys330^ by introducing the following mutations: C239L, C249S, C426T, C436V, and C443S. Since C0-C2^Cys249^ contains an endogenous cysteine at position 249, the remaining four endogenous cysteines were similarly removed. The following N-terminal cMyBP-C HCM mutations were introduced to each C0-C2^S85C^, C0-C2^Cys249^, and C0-C2^P330C^ construct: T59A, R282W, E334K, L349R, and L352P. The results were C0-C2 constructs containing a single cysteine in the C0, C1, or M-domain with an cMyBP-C HCM mutation inserted: C0-C2^S85C, T59A^, C0-C2^S85C, R282W^, C0-C2^S85C, E334K^, C0-C2^S85C, L349R^, C0-C2^S85C, L352P^, C0-C2^Cys249, T59A^, C0-C2^Cys249, R282W^, C0-C2^Cys249, E334K^, C0-C2^Cys249, L349R^, C0-C2^Cys249, L352P^, C0-C2^P330C, T59A^, C0-C2^P330C, R282W^, C0-C2^P330C, E334K^, C0-C2^P330C, L349R^, C0-C2^P330C, L352P^. All sequences were confirmed by DNA sequencing (Eton Biosciences). Protein production in E. coli BL21DE3-competent cells (New England Bio Labs) and purification of C0-C2 protein using His60 Ni Superflow resin were done as described (24). C0-C2 (with His-tag removed by TEV protease digestion) was further purified using size-exclusion chromatography to achieve >90% intact C0-C2 as described (36) and then concentrated, dialyzed to 50/50 buffer (50 mM NaCl and 50 mM Tris, pH 6.7), and stored at 4°C.

C0-C2 was labeled with tetramethylrhodamine (TMR) as the acceptor probe in 50/50 buffer in dim lighting conditions. C0-C2 (50 μM) was first treated with the reducing agent TCEP (200 μM) for 30 min at 23°C while rocking. TMR was added (stored as 20 mM stock in DMF at -80°C) to a final concentration of ∼100 μM (concentration of dye is dependent on the accessibility of the cysteine for labeling and may need to be adjusted accordingly). Labeling was done for 1 h at 23°C and terminated by the addition of 5x molar excess of DTT. Unincorporated dye was removed by extensive dialysis against the final buffer respective to myosin or actin TR-FRET experiments. Following the dialysis, labeled C0-C2 was centrifuged for 30 min at 100,000 RPM (350,000 *xg*) in a Beckman TLA-120.2 rotor to remove any precipitated dye and insoluble protein. The degree of labeling would range from 50-90% dye/C0-C2 as measured by UV-vis absorbance and by TMR’s extinction coefficient. To attain 50-60% dye/C0-C2 for TR-FRET experiments, the unlabeled corresponding C0-C2 construct was added.

### Protein and Dye Concentration

The Bradford protein concentration assay (BCA) and SDS-Page Gel using a known a known BSA protein standard was used throughout this study to determine protein concentrations. The extinction coefficient for FMAL is 68,000 at 494 nm and the extinction coefficient for TMR is 91,000 at 542 nm, provided by manufacturer’s specifications.

### In vitro phosphorylation of cMyBP-C

C0-C2 was treated with 7.5 ng PKA/μg C0-C2 at 30°C for 30 min. This is 3x the level (2.5 ng PKA/μg C0-C2) needed to achieve maximal phosphorylation as determined by in-gel staining of proteins with Pro-Q Diamond (24, 36).

### Time-Resolved FRET (TR-FRET) data acquisition and analysis

Fluorescence lifetime measurements were acquired using a high-precision fluorescence lifetime plate reader (FLTPR; Fluorescence Innovations, Inc) (23, 24, 37), provided by Photonic Pharma LLC. For TR-FRET experiments, FMAL was excited with a 473-nm microchip laser (Bright Solutions) and emission was filtered with 488-nm long-pass and 517/20-nm band-pass filters (Semrock). The photomultiplier tube (PMT) voltage was adjusted so that the peak signals of the instrument response function (IRF) and the biosensor were similar. The observed waveforms were convolved with the IRF to determine the lifetime (τ) (Eq. 1) by fitting to a single-exponential decay (23, 24, 37). The decay of the excited state of the fluorescence FMAL dye attached to myosin at V105C or actin at Cys-374 to the ground state is:

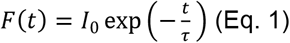

where *I*_0_ is the fluorescence intensity upon peak excitation (*t* = 0), and τ is the fluorescence lifetime (*t* = τ when *I* decays to 1/*e* or ∼37% of *I*_0_). The efficiency of energy transfer E (FRET Efficiency) was calculated from the average time of the donor in presence (τ_DA_) and absence (τ_D_) of acceptor (Eq. 2):

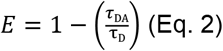

### Determination of FRET_max_ and Kd values

The maximum FRET (FRET_max_) and K_d_ values for C0–C2 binding to myosin and actin were determined by fitting the data to a quadratic model (Michaelis–Menten function) using Origin Pro 2019 computer software package through a nonlinear least-squares minimization (Levenberg– Marquardt algorithm) as previously described in Bunch *et al*. (24, 34). These apparent K_d_ and FRET_max_ values are used as comparative indicators of binding characteristics for C0-C2 binding to myosin and actin under different conditions (± phosphorylation or in the presence of mutations). The adjusted R^2^ was used to identify the goodness-of-fit and supports whether the K_d_ and FRET_max_ generated had meaning. In TR-FRET generated curves, the K_d_ values represent the apparent dissociation constants (C0-C2 concentration required for half-maximal binding).

### Determination of *Z′*-Factor for C0-C2^Cys249^

For suitability in high-throughput screening (HTS), TR-FRET assay quality was determined for unphosphorylated versus phosphorylated donor-acceptor labeled C0-C2^Cys249^ in the presence of donor-labeled myosin or actin. The lifetimes of unphosphorylated donor-acceptor labeled C0-C2 ^Cys249^ (τ _A_) and phosphorylated donor-acceptor labeled C0-C2 ^Cys249^ (τ_B_) were compared and indexed by the *Z′-*factor (Eq.3):

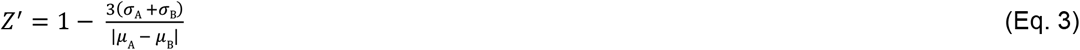

where *σ*_A_ and *σ*_B_ are the standard deviations (S.D) of the τ_A_ and τ_B_ lifetimes, respectively, *μ*_A_ and *μ*_B_ are the means of the τ_A_ and τ_B_ lifetimes, respectively. *Z′*-factor of less than 0 is “useless”, 0 to 0.5 is “good” and 0.5 to 1.0 is “excellent” assay quality (28).

### Statistics

Average data are provided as mean ± standard error (SE). Each experiment was done with >2 separate protein preparations. Statistical significance (*p*<0.05) is evaluated by use of Student’s t-test. Biological repeats (N, independent protein preparations) and technical repeats (n, independent reactions) are as indicated in each figure and table.

## Acknowledgments

This work was supported by NIH grants R01 HL141564 (to B.A.C.) and T32 HL007249 (to C. Gregorio and J. Konhilas, University of Arizona)

## Author Contributions

F.L.W., T. A. B., and B. A. C. conceptualization; F.L.W., T. A. B., V. C. L., and B. A. C. formal analysis; F.L.W., T. A. B., and B. A. C. supervision; F.L.W., T. A. B., V. C. L., and B. A. C. validation; F.L.W., T. A. B., V. C. L., and B. A. C. investigation; F.L.W., T. A. B., and B. A. C. visualization; F.L.W., T. A. B., V. C. L., and B. A. C. methodology; F.L.W., T. A. B., and B. A. C. writing-original draft; F.L.W., T. A. B., V. C. L., and B. A. C. writing-review and editing; V. C. L. and B. A. C. project administration; F.L.W. and B. A. C. funding acquisition; B. A. C. resources.

## Competing Interest Statement

B.A.C. serves as President of BC Biologics LLC. This relationship has been reviewed and managed by the University of Arizona. BC Biologics had no role in this study. B.A. Colson and F.L. Wong filed a provisional patent application (patent pending, application no. 63/076,735) and B.A. Colson filed a patent application (patent pending, application no. PCT/US21/14142) based on this work. The other authors declare no competing financial interests.

