## Supplemental Figures and Table for "N-terminal cardiac myosin-binding protein C interactions with myosin and actin filaments using time-resolved FRET"

### Supplementary Figures and Table

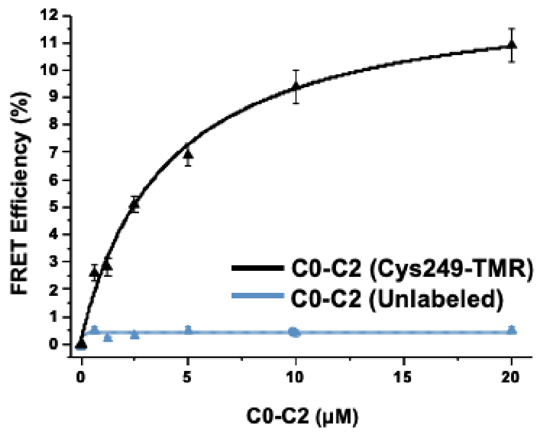


**Fig. S1.** To calculate FRET Efficiency for myosin TR-FRET, the donor-only sample consists of 1 μM of myosin and buffer rather than 1 μM of myosin and a respective concentration of unlabeled C0-C2. The FRET Efficiency of adding unlabeled C0-C2 remained <1%, therefore the use of buffer for the donor-only measurement is more practical in sparing protein and produces the same results.

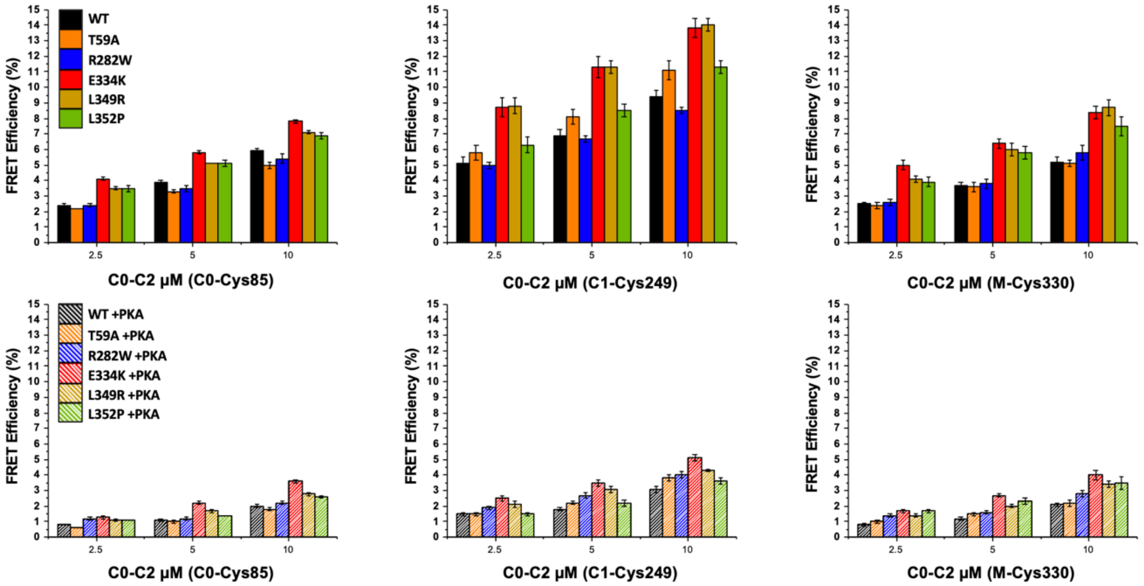


**Fig. S2.** Myosin TR-FRET testing of HCM mutants inserted in C0-C2 with probes located in the C0, C1, or the M-domain. Three concentrations of C0-C2 were tested (2.5, 5, 10 µM). Unphosphorylated C0-C2 is shown above in solid bars. Phosphorylated C0-C2 is shown below in stripped bars. WT is shown in black. HCM mutants T59A (orange), R282W (blue), E334K (red), L349R (gold), and L352P (green) were tested. Data are provided as mean ± SE (N=2-3, n=10-15).

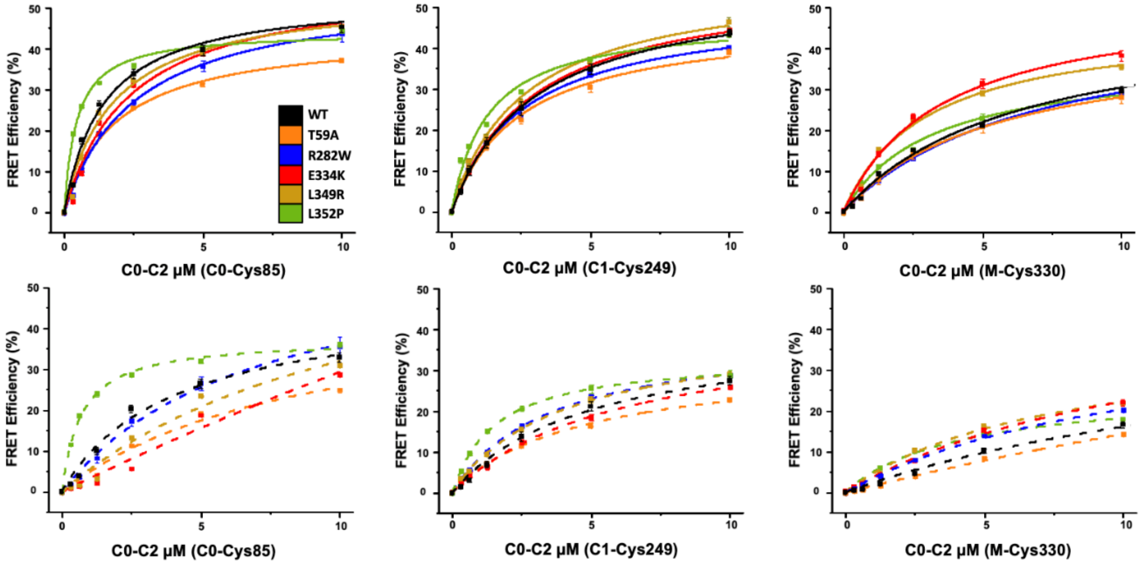


**Fig. S3.** Actin TR-FRET testing of HCM mutants inserted in C0-C2 with probes located in the C0, C1, or the M-domain. Due to the feasibility of actin TR-FRET compared to myosin TR-FRET, a range of concentrations were tested to generate binding curves. Unphosphorylated C0-C2 is shown above in solid lines. Phosphorylated C0-C2 is shown below in dotted lines. WT is shown in black. HCM mutants T59A (orange), R282W (blue), E334K (red), L349R (gold), and L352P (green) were tested. Data are provided as mean ± SE (N=2, n=8-10).

**Table S1. TR-FRET Binding parameters of cMyBP-C C0-C2 binding to myosin and actin.**

| | Probe Site | PKA | $K_D$ | $FRET_{Max}$ | Adj. $R^2$ |
|---|---|---|---|---|---|
| Myosin | C0-Cys85 | - | 9.0 ± 2.6 | 11.6 ± 1.6 | 0.96 |
| | | + | 125.8 ± 314.6 | 29.5 ± 65.4 | 0.88 |
| | C1-Cys249 | - | 3.4 ± 1.1 | 12.5 ± 1.4 | 0.93 |
| | | + | 13.2 ± 12.6 | 8.3 ± 4.2 | 0.60 |
| | M-Cys330 | - | 6.9 ± 2.5 | 12.6 ± 2.0 | 0.93 |
| | | + | 9.7 ± 5.3 | 16.1 ± 15.7 | 0.65 |
| Actin | Probe Site | PKA | $K_D$ | $FRET_{Max}$ | Adj. $R^2$ |
| | C0-Cys85 | - | 1.4 ± 0.2 | 53.2 ± 1.6 | 0.99 |
| | | + | 4.1 ± 0.6 | 47.1 ± 2.6 | 0.99 |
| | C1-Cys249 | - | 3.0 ± 0.1 | 55.9 ± 0.5 | 1.00 |
| | | + | 5.0 ± 0.5 | 40.8 ± 1.7 | 1.00 |
| | M-Cys330 | - | 6.3 ± 0.7 | 49.4 ± 2.2 | 0.99 |
| | | + | 17.8 ± 2.8 | 44.3 ± 4.0 | 1.00 |

A range of C0-C2 concentrations (1.25 to 20 µM) with acceptor sites in C0 (S85C), C1 (Cys249), and M-domain (P330C) (-/+ PKA phosphorylation) were tested with 1 µM of myosin or actin by TR-FRET. Data were fit to a quadratic model (Michaelis-Menten function) to generate $K_d$, $FRET_{Max}$, and $R^2$ values (see Materials and Methods). Red numbers indicate binding curves with an Adj. $R^2$ of <0.89.
